# Characterisation of subpopulations of chicken mononuclear phagocytes that express TIM4 and the macrophage colony-stimulating factor receptor (CSF1R)

**DOI:** 10.1101/294504

**Authors:** Tuanjun Hu, Zhiguang wu, Stephen J. Bush, Lucy Freem, Lonneke Vervelde, Kim M. Summers, Adam Balic, David A. Hume, Pete Kaiser

## Abstract

The phosphatidylserine receptor, TIM4, encoded by *TIMD4*, mediates the phagocytic uptake of apoptotic cells. We applied anti-chicken TIM4 monoclonal antibodies, in combination with *CSF1R* reporter transgenes to dissect the function of TIM4 in chick (*Gallus gallus*). During development *in ovo*, TIM4 was present on the large majority of macrophages but expression became more heterogeneous post-hatch. Blood monocytes expressed KUL01, class II MHC and *CSF1R-*mApple uniformly. Around 50% of monocytes were positive for surface TIM4. They also expressed many other monocyte-specific transcripts at a higher level than TIM4^−^ monocytes. In liver, highly-phagocytic TIM4^hi^ cells shared many transcripts with mammalian Kupffer cells and were associated with uptake of apoptotic cells. Although they expressed *CSF1R* mRNA, Kupffer cells did not express the *CSF1R*-mApple transgene, suggesting that additional *CSF1R* transcriptional regulatory elements are required by these cells. By contrast, *CSF1R*-mApple was detected in liver TIM4^lo^ and TIM4^−^ cells which were not phagocytic and were more abundant than Kupffer cells. These cells expressed *CSF1R,* alongside high levels of *FLT3*, *MHCII, XCR1* and other markers associated with conventional dendritic cells (cDC) in mice. In bursa, TIM4 was present on the cell surface of two populations. Like Kupffer cells, bursal TIM4^hi^ phagocytes co-expressed many receptors involved in apoptotic cell recognition. TIM4^lo^ cells appear to be a sub-population of bursal B cells. In overview, TIM4 is associated with phagocytes that eliminate apoptotic cells in the chick. In the liver, TIM4 and *CSF1R* reporters distinguished Kupffer cells from an abundant population of DC-like cells.

## Introduction

Phagocytosis of apoptotic or senescent cells by macrophages is a physiological process for maintenance of cell populations in tissues during embryonic development and adult homeostasis (1, 2). Apoptotic cells are recognized by phagocytes through multiple mechanisms. One depends upon the exposure of the normal inward-facing phosphatidylserine (PS) of the lipid bilayer to the outer layers of the plasma membrane (3). T cell immunoglobulin and mucin domain-containing 4 (TIM4), encoded by the *TIMD4* locus, was defined as a plasma membrane PS receptor, (4). *Timd4* in mice is expressed primarily by subsets of macrophage lineage cells in a restricted set of tissues, notably Kupffer cells in the liver, which mediate clearance of senescent red blood cells (5). Resident mouse peritoneal macrophages also express high levels of TIM4 which is essential for their recognition of apoptotic cells. However, in other locations where *Timd4* is highly-expressed, such as the marginal zone in spleen, TIM4 is not essential for apoptotic cell recognition (6). In mouse liver, *Timd4* provides a marker for macrophages of embryonic origin, that reside together with, but are distinct from, those recruited from blood monocytes (5). Deficiency of *Timd4* in mice produces T and B cell hyperactivity and autoimmunity, attributed to the failure to regulate antigen-reactive T cell differentiation (7). Unlike other TIM family members, TIM4 has no tyrosine kinase motif in its cytoplasmic tail (8). Accordingly, other PS receptors or co-receptors in addition to TIM4 are required to initiate particle uptake and signal transduction. Recognition of PS by TIM4 may also contribute to macropinocytosis of viruses (9, 10) notably in association with TIM1, encoded by the adjacent *Havcr1* locus.

We previously identified the chicken *TIMD4* locus, and produced monoclonal antibodies against two distinct isoforms of the TIM4 protein (11). Recombinant chicken TIM4 bound to PS, and like its mammalian orthologue, is thereby implicated in recognition of apoptotic cells. A TIM4-fusion protein also had co-stimulatory activity on chicken T cells, suggesting a function in antigen presentation (11). In birds, as in mammals, macrophage differentiation depends upon signals from the CSF1 receptor (CSF1R), which has two ligands, CSF1 and IL34 (12). In contrast to the mammalian system, in chickens TIM4 was highly-expressed by macrophages grown *in vitro* in macrophage colony-stimulating factor (CSF1).

Anti-CSF1R antibodies (13) and transgenic reporter genes based upon control elements of the *CSF1R* locus (14) provide convenient markers for cells of the macrophage lineage in birds. An emerging view in mammalian macrophage development is that many tissue macrophage populations are maintained by self-renewal of macrophages seeded from yolk sac-derived progenitors during embryonic development, independently of blood monocytes (15, 16). This is less evident in chickens, where intra-embryonic transplantation of bone marrow precursors gave rise to donor-derived macrophages throughout the body (17). Nevertheless, the first evidence that macrophages are produced by the yolk sac derived from studies of chicken development and these cells are involved extensively in the clearance of apoptotic cells (reviewed in (18)). A recent study of the time course of chicken embryonic development based upon cap analysis of gene expression (CAGE) detected expression of both *TIMD4* and *HAVCR1* (encoding the related TIM1 protein) around day 2 of development, when the first CSF1R-dependent macrophages are also detected (19, 20). In the current study, we utilise anti-TIM4 antibodies in combination with a *CSF1R-*mApple reporter to locate and characterise the gene expression profiles of cells that express this receptor. We show that TIM4^+^ Kupffer cells in chicken closely resemble mammalian Kupffer cells and are distinct from TIM4^+^ phagocytes in the bursa. *Inter alia*, our analysis led to the identification of an abundant population of TIM4^lo/-^ *CSF1R*-mApple^+^ cells in the liver that resemble mammalian antigen-presenting dendritic cells but are much more abundant that their counterparts in mouse liver. Their identification and location suggests that the avian liver has a function in the control of T cell-mediated immune responses.

## Materials and Methods

### Chicken and antibodies

The J line chicken was cross-bred from nine lines, originally inbred from Brown Leghorn chickens at the Poultry Research Centre, Edinburgh to study a variety of traits, such as egg laying, plumage and vigour (http://www.narf.ac.uk/chickens/lines). This strain of chicken expressed multiple TIM4 isoforms (11). *CSF1R*-mApple transgenic chickens, which carry the chicken *CSF1R* regulatory sequences directing expression of the red fluorescent protein mApple to the cytoplasm of macrophages (14) and commercial Novogen Brown layers were also included in this study. All birds were hatched and housed in premises licensed under a UK Home Office Establishment License in full compliance with the Animals (Scientific Procedures) Act 1986 and the Code of Practice for Housing and Care of Animals Bred, Supplied or Used for Scientific Purposes. All procedures were conducted under Home Office project licence PPL 70/7860, according to the requirements of the Animal (Scientific Procedures) Act 1986, with the approval of local ethical review committees. Animals were humanely culled in accordance with Schedule 1 of the Animals (Scientific Procedures) Act 1986.

Monoclonal antibody JH9 against chicken TIM4-extracellular-domain (amino acids 1-209) was raised and characterised as described previously (11). JH9 was labelled with AlexaFluor-647 for flow cytometric analysis, or with AlexaFluor-568 (Invitrogen, Paisley, UK) for immunofluorescent microscopy analysis, as per manufacturer’s instructions. Anti-chicken CSF1R (ROS-AV170) (13) was also labelled by AlexaFluor-568 (Invitrogen) for immunofluorescent microscopy analysis. Other primary antibodies included anti-chicken CD45-FITC (clone UM16-6, Bio-Rad), FITC-labelled anti-chicken monocyte/macrophage marker KUL01 (clone KUL01, Southern Biotech), anti-Bu-1-FITC (clone AV20, Southern Biotech), anti-chicken MHC II-FITC (2G11, Southern Biotec), IgG1 isotype control, mouse anti-ovine CD335 (GR13.1), rabbit anti-GFP (AF-688-labelled) (ThermoFisher, UK), rabbit anti-RFP (DsRed) (BioVision, CA, USA; distributed by Cambridge Bioscience, UK.

### Isolation and culture of chicken primary cells

Kupffer cells were isolated from livers dissected from 4-6 week old Novogen Brown layers, based upon a method previously developed for the mouse (21). The liver was chopped into small pieces, transferred into 10 ml of 1 mg dispase/collagenase D (Roche Applied Scientific) and digested by incubation at 37 ˚C for 30 min with occasional gentle mixing. The tissue was passed through a 100 µm cell strainer and the cell suspension was centrifuged at 300× *g* for 5 min at 4 ˚C. The pellet was washed twice with 50 ml RPMI 1640 then centrifuged at 50× *g* without the brake for 3 min at 4 °C to sediment parenchymal cells. This process was repeated. To remove cell debris and red blood cells, the non-parenchymal cell pellet was resuspended in 5 ml RPMI 1640, gently overlaid onto the same volume of Histopaque 1.077 and centrifuged at 400 × *g* without brake for 30 min. The cells at the interface between RPMI medium and Histopaque were carefully collected and washed twice with RPMI 1640 medium.

Blood leukocytes were isolated from 4-8 week old *CSF1R*-mApple transgenic chickens (14) by density gradient centrifugation using lymphoprep (density 1.077± 0.001g/ml, Alere Technologies, Norway) as described previously (22). Non-parenchymal cells in liver tissues from 4-8 week old *CSF1R*-mApple transgenic chickens were prepared as described above. A similar digestive procedure using dispase/collagenase D was also applied to bursal tissues from *CSF1R*-mApple transgenic chickens; the tissue digest was passed through 70 µm cell strainer for single bursal cell suspension.

### Phagocytosis assays

For analysis *in vitro*, isolated Kupffer cells were seeded into 4-well glass chamber slides (Nunc) and cultured in Dulbecco’s Modified Eagle’s Medium (DMEM) supplemented with 10% fetal bovine serum, 1× glutamax and 100 U/ml penicillin/streptomycin at 41 °C in a 5% CO_2_ incubator overnight. Non-adherent cells were then removed from the wells. To test the phagocytic activity of the adherent Kupffer cells, Zymosan A Bioparticles (Thermo Scientific) or apoptotic chicken thymocytes were diluted in DMEM and added to the wells at approximately 10 particles/cell. The cells were then incubated at 41°C for 2 h, washed four times with ice-cold PBS, fixed with 4% paraformaldehyde (PFA) for 20 min and permeabilized by 1% Triton X-100 for 15 min. The cells were probed by AlexaFluor-568-conjugated anti-TIM4 monoclonal antibody JH9 and the nucleus was counterstained with DAPI. Images were taken using a Leica DMLB microscope. Chicken red blood cells were isolated from chicken whole blood by Histopaque (1.077) gradient and aged by incubation at 4°C for thirty hours. They were added to adherent Kupffer cells at a ratio of 10:1. The cells were incubated at 41°C overnight. As above, the cells were then washed, fixed and stained with haematoxylin (Sigma-Aldrich) for 2 min. The results were analysed using a Nikon Ni-E microscope.

To confirm phagocytic activity *in vivo*, 4 week old *CSF1R*-mApple transgenic chickens were injected intravenously with 100μl of 0.1μ diameter FluoSpheres^®^ (ThermoFisher). Birds were culled 3 hours after administration of beads by cervical dislocation and tissues were removed and fixed in 4% PFA. Fixed tissues were processed for immunostaining as detailed below.

### Flow-cytometry

Single cell suspensions from embryos, blood, liver or bursa were used for flow cytometry analysis. Cells were washed with FACS buffer (PBS, 0.5% bovine serum albumin (BSA) and 0.05% sodium azide) and incubated with anti-TIM4-AF647 and FITC-conjugated antibody to other markers including CD45, KUL01, MHC class II or Bu-1 as described above. Cells were incubated at 4°C in the dark for 30 min and washed three times in FACS buffer. Cells were resuspended in 300 μl PBS with SYTOX® Blue Dead Cell Stain (1.0 μM, Invitrogen) 5 min prior to analysis using a BD LRSFortessa (BD Biosciences, UK). At least 100,000 events were acquired. Dead cells were excluded by SYTOX® Blue staining and doublets were discriminated based on signal processing (SSC-A/H or FSC-A/H). Data were analysed using FlowJo software (FlowJo, Ashlan, OR. USA).

### Separation and gene expression profiling by RNAseq

Blood leukocytes or isolated cells from liver and bursa were purified into different populations using BD FACS Aria IIIu, based on their expression of TIM4, *CSF1R-*mApple or Bu-1. Blood leukocytes and non-parenchymal liver cells from *CSF1R-*mApple reporter birds were labelled using anti-TIM4 (AF647-labelled) or bursal cells by anti-TIM4 and Bu-1 (B cell surface marker) for 30 min at 4 °C and separated by FACS. All gate settings were based on isotype-matched controls. The separated cell populations (TIM4^+^mApple^+^ and TIM4^−^mApple^+^ from blood; TIM4^+^mApple^−^, TIM4^+^mApple^+^ and TIM4^−^mApple^+^ from liver; TIM4^+^ Bu-1^−^, TIM4^+^ Bu-1^+^ and TIM4^−^ Bu-1^+^ from bursa) were then lysed with TRIzol reagent (Invitrogen). RNA was extracted using RNeasy mini kit (Qiagen). RNA quality (RIN > 7) was assessed by Agilent RNA screen tape assay with Agilent 2200 TapeStation and quantified by a Qubit RNA HS kit (Molecular Probes). cDNA libraries were prepared using NEBNext Ultra RNA Library Prep kit for Illumina (New England BioLabs), with each sample containing different index primer (NEBNext Multiplex Oligos for Illumina, Index primer Set 1). Pools of 4 libraries were sequenced by Edinburgh Genomics using the Illumina HiSeq 4000 Sequencing System. Bone marrow-derived macrophages were prepared by cultivation of bone marrow cells in macrophage colony-stimulating factor (CSF1) as described (12, 17). mRNA was isolated and gene expression profiles assayed as above. For comparative analysis, bursa and spleen mRNA was prepared as above from 8-12 newly-hatched birds as part of a separate project looking at gender and genotype-associations with gene expression profiles (Ms in preparation). For the current analysis, the expression of each transcript was averaged across the whole data set. All of the sequencing data generated for this project is deposited in the European Nucleotide Archive (ENA) under study accession number PRJEB25788 (http://www.ebi.ac.uk/ena/data/view/PRJEB25788).

### Differential expression analysis

Expression level was quantified, as both transcripts per million (TPM) and estimated read counts, using Kallisto v0.42.4 (23). Transcript-level read counts were summarised to the gene level using the R/Bioconductor package tximport v1.0.3 (24) with gene names obtained from the GalGal5.0 annotation (via Ensembl BioMart v90). The tximport package aggregates Kallisto output into a count matrix, useable by the R/Bioconductor package edgeR v3.14.0. for differential expression analysis, as well as calculating an offset that corrects for changes to the average transcript length between samples (which can reflect differential isoform usage). Using edgeR, gene counts were normalised using the ‘trimmed mean of M values’ (TMM) method, with a negative binomial generalized log-linear model fitted, and p-values corrected for multiple testing according to a false discovery rate (FDR).

### Immunostaining of tissue sections

Tissue slices were fixed in 4% paraformaldehyde (PFA)/PBS for 1 h, washed with PBS, equilibrated with 15% sucrose/PBS overnight at 4°C, embedded in OCT compound and flash frozen in liquid nitrogen. Tissue sections (10 µm) were cut, mounted onto sugar-coated slides and dried overnight. For immunohistochemistry, sections were rehydrated with PBS for 5 mins; endogenous peroxidase activity was quenched by incubating slides with 0.3% H_2_O_2_ in PBS. After blocking with normal horse serum, primary antibodies were added to sections and incubated at 4˚C overnight, followed by secondary antibody biotinylated goat-anti-mouse IgG for 30 min then avidin-biotin-peroxidase complex (GE Healthcare, UK). Vector AEC (Sigma) or NovaDAB substrate was added to reveal bound peroxidase, and sections were counterstained with haematoxylin. The resultant staining was analysed using a Nikon Ni microscope. For immunofluorescent staining, the sections were rehydrated and blocked with horse serum (14) the fluorescence-labelled antibodies were added to sections and incubated overnight. The resulting staining was analysed using a Leica DMLB microscope.

For whole-mount TIM4 staining of embryonic tissues, 4% PFA fixed tissues were placed in PBS with 10% normal horse serum, 0.1% Triton-X100 for two hours at 4˚C on a rocking platform, followed by overnight incubation with anti-TIM4 antibody. Tissues were washed for 30 minutes in PBS, incubated for two hours with donkey anti-mouse AlexaFluor-647 (Invitrogen, Paisley, UK) secondary antibody and washed again for 30 minutes before imaging. For imaging of whole mount immunostained tissues, a 35 mm × 10 mm petri dish was modified by cutting a hole in the lower section and fixing a coverslip over the hole with nail polish. Stained embryonic limb buds were placed on the coverslip, with the lower surface in contact with the cover slip. Samples were imaged using an inverted confocal microscope (Zeiss LSM710). For 3D-rendering, confocal z-stacks were created by obtaining images at 0.45 μm intervals. Images were captured using Zeiss ZEN software and analysed using Imaris software, version 8.2 (Bitplane).

### Terminal deoxynucleotidyl transferase (TdT) dUTP Nick-End Labeling (TUNEL) staining

TUNEL staining was carried out as previously (25) with minor modifications. After rehydration sections were incubated with 20% FCS and 1% BSA in PBS for 1□hour. For nick-end labelling, the sections were equilibrated by incubation in terminal deoxynucleotidyl transferase (TdT) buffer (0.2□M potassium cacodylate, 1□mM cobalt chloride, 25□mM Tris and 0.01% Triton X-100) for 15□min. Labelling reaction mixture (10 U TdT (ThermoFisher) and 1 nM biotinylated dUTP (Roche) in TdT buffer) was added to sections and incubated for 1□hour at 37□°C in a humidified chamber. The labelling reaction was terminated by washing the slides three times in TB buffer (sodium chloride 300 mM, sodium citrate 30 mM). To visualize TUNEL labelled cells, sections were incubated with FITC conjugated streptavidin (ThermoFisher) for 1 h. Sections were then stained with Alexa Fluor-568-conjugated anti-TIM4 for 2 h and the nucleus was stained with DAPI. The resulting images were visualised using a Leica DMLB microscope.

### Western blot

Bursa of Fabricius, spleen and liver were taken from 6 week old birds. Tissues were disrupted using lysing matrix D (MP Biomedical, Loughborough Leicester, UK) with lysis buffer (1% NP40/PBS) in a Fastprep 24 homogeniser (MP Biomedical); the lysates were centrifuged at 13,000× *g* for 5 min to remove any cell debris and DNA. 30 µg of lysate protein was separated in 4-15% SDS-PAGE (Bio-rad, Hertfordshire, UK). After trans-blotting protein onto PVDF membrane (Sigma-Aldrich), the membrane was probed with primary antibodies at 4°C overnight, followed by horse radish peroxidase (HRP)-conjugated secondary antibody for 2 h at room temperature. HRP activity was detected using Enhance Chemiluminescent (ECL) substrate (ThermoFisher).

## Results

### Distribution of TIM4-positive cells in chickens post hatch

*TIMD4* mRNA was previously found to be relatively abundant in the majority of non-immune as well as immune-related tissues of the chicken (11). Although macrophages are a major component of all tissues, highlighted using the *CSF1R-*reporter gene (14) this observation suggests either that the expression of *TIMD4* by macrophages is much less tissue specific than in mammals, or else that *TIMD4* is expressed by non-macrophage lineage cells. To distinguish these alternatives, sections of tissues from 6 week old chicks were examined by immunohistochemistry (IHC) using KUL01 (monocyte/macrophage marker) and anti-chicken TIM4 antibody JH9 (11).

To validate the monoclonal antibody binding, tissue extracts were first analysed by western blot. As shown in Figure 1A, in bursa of Fabricius, the monoclonal antibody JH9 bound to a single TIM4 product at 100 kDa, the predicted size encoded by the long isoform TIM4L_1_ mRNA (11). In spleen, two TIM4 products at the similar density were detected, the larger band at 150 kDa predicted by the longer TIM4L_0_ mRNA (11). In liver, the 150 kDa isoform was most highly-expressed, but a minor larger isoform was also detected. Figure 1B shows the patterns of staining of TIM4 in a range of adult tissues, compared to the widely-used macrophage marker, KUL01, which is encoded by a the likely orthologue (*MRC1-LB*) of the mammalian mannose receptor gene, *MRC1* (26), itself regarded as a marker of functional polarization in mammalian macrophages (27). In thymus TIM4^+^ / KUL01^+^ cells were mainly located in the medulla (Figure 1B). In the caecal tonsil, an important gut-associated lymphoid tissue (GALT), numerous stellate KUL01 and TIM4-positive cells were scattered throughout the lymphoid tissue within the lamina propria and the submucosa (arrow). Fewer KUL01 and TIM4-positive cells were also present in the muscularis mucosae (M) (Figure 1B). In lung, both antibodies labelled numerous cells within the lung parenchyma and the bronchial walls (Figure 1B). In the bursa of Fabricius, staining patterns with the two antibodies were quite distinct. Whereas KUL01 strongly stained interfollicular cells and weakly stained the cells clustered at corticomedullary epithelium, most TIM4^+^ cells were scattered within bursal lymphoid follicles, particularly at the corticomedullary epithelium (Figure 1B). In spleen, KUL01 and TIM4^+^ cells were mostly distributed at the interface between peri-ellipsoidal white pulp (PWP) and red pulp, a region equivalent to the mammalian marginal zone, and fewer KUL01 and TIM4-positive cells were dispersed in red pulp (RP) (Figure 1B). In small intestine (jejunum) both KUL01 and anti-TIM4 stained presumptive macrophages in the lamina propria of the villi and crypts (Figure 1B). In the liver, KUL01 and TIM4-positive cells were clearly separate and morphologically distinct. KUL01^+^ cells were relatively small and round where TIM4^+^ cells were stellate and irregular, lining the walls of hepatic sinusoids, consistent with their identity as Kupffer cells (Figure 1B). In testis, KUL01^+^ and TIM4^+^ cells appeared similar, located in the connective tissue surrounding seminiferous tubules and within aggregates of lymphoid cells (Figure 1B). In overview, TIM4 was much more widely-expressed in chickens than in mice, apparently restricted to macrophage-like cells, but not completely coincident with the widely-used KUL01 marker.

**Figure 1.**
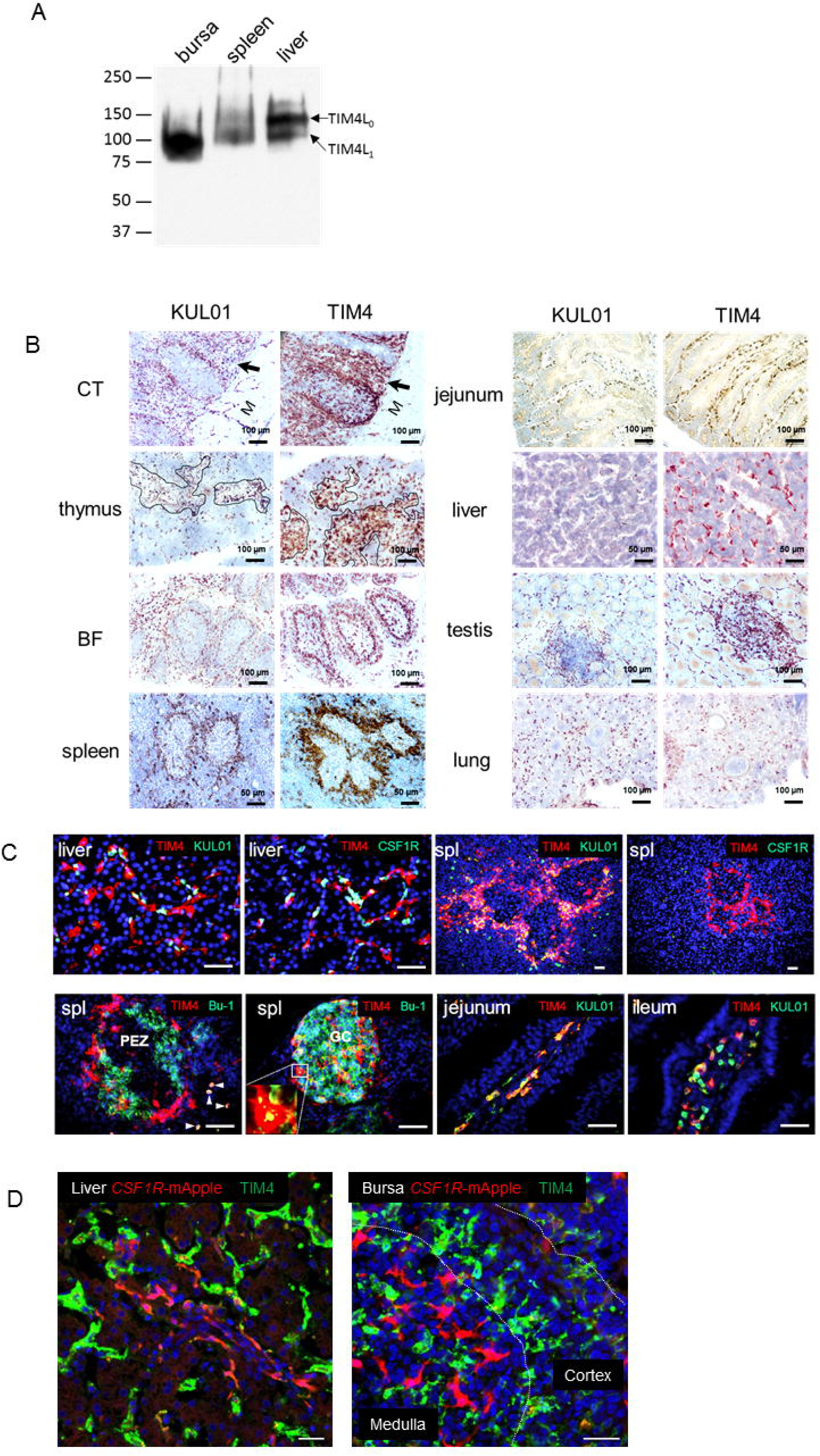
A). Western blot analysis of TIM4 protein expression. Bursa, spleen and liver were isolated from 6 week old birds. The protein products of TIM4 isoforms, indicated by arrows, in tissue lysates were detected using JH9 antibody. The protein marker sizes are in kDa. The experiment was repeated twice. B). Immunohistochemistry of TIM4 macrophages in peripheral tissues. Cryosections from 6-week-old birds were stained for TIM4 or KUL01 as indicated. Positive staining was detected by DAB substrate (brown) or AEC (red, in liver) and nuclei were counterstained with hematoxylin (blue). The follicles in thymus are outlined. BF: bursa of Fabricius, CT: caecal tonsil (M in this section marks the muscularis). The results show representative staining of serial sections from at least three birds. C) Heterogeneity of resident TIM4 macrophages. Cryostat sections (10 µm) from 4 week old birds were co-stained with anti-TIM4 (red) or KUL01, CSF1R or Bu-1 (green) as indicated. The tissues included liver, spleen (spl), jejunum and ileum. PEZ: Splenic peri-ellipsoid zone. Arrows highlight TIM4^+^ Bu-1^+^ cells. GC: splenic germinal centre (GC). The inserted picture highlights a TIM4^+^ macrophage resembling a mammalian tingible body macrophage, closely-associated with B cell debris. D) Cryostat sections (10 µm) of liver and bursa of Fabricius from 4-week-old *CSF1R*-mApple transgenic birds stained for TIM4 (green). The nucleus was counterstained by DAPI (blue). Scale bar, 50 µm. The images represent one of three serial sections from at least three birds.

Figure 1C examines the heterogeneity of TIM4 expression by immunofluorescence, with antibodies against CSF1R (13) and Bu-1 (expressed by subsets of macrophages as well as B cells (28)) as additional markers. Consistent with the IHC data in Figure 1B, TIM4^+^ and KUL01 appeared mutually exclusive in liver. By contrast, in spleen, peri-ellipsoidal macrophages co-expressed TIM4 and KUL01. CSF1R protein was not detectable by immunofluorescence on any TIM4^+^ cells. Co-staining of spleen sections using anti-TIM4 and Bu-1 antibodies highlighted the architecture of the peri-ellipsoid zone in spleen surrounded by TIM4^+^ macrophages. TIM4^+^ Bu-1^+^ cells were also evident scattered in red pulp. The staining of spleen revealed the structure of the B cell-rich germinal centre, in which TIM4^+^ macrophages not only outlined the germinal centre, structurally similar to mammalian metallophilic macrophages, but also intermingled with B cells inside germinal centre to engulf B cell debris, equivalent to tingible-body macrophages in mammalian germinal centres.

In the intestine, the staining of jejunum and ileum indicated that a heterogeneous population of TIM4^+^, KUL01^+^ and TIM4^+^ KUL01^−^ cells localized at the lamina propria of the villi. In the liver, the *CSF1R-*mApple transgene was detectable on only a small subset of cells, as previously noted (14). By contrast, TIM4 brightly labelled an almost continuous network of cells (Figure 1D), consistent with the high levels of TIM4 protein detected on Western Blots. In the bursa of Fabricius, *CSF1R-*mApple positive cells formed an extensive network of stellate cells within the medulla, but again there was little apparent overlap with TIM4 (Figure 1D). These results indicate that in chickens, as in mice, TIM4 is a unique marker for Kupffer cells and subpopulations of macrophages in lymphoid organs. By contrast to mice, TIM4 is widely-expressed by resident macrophage-like cells throughout adult tissues in the chicken, whereas niether the *CSF1R-*mApple transgene, nor KUL01, were uniformly expressed by all chicken tissue macrophages.

### Phagocytic activity of TIM4 macrophage

Because they are the largest macrophage population with direct contact to the blood, Kupffer cells contribute to the clearance of the damaged or aging erythrocytes as well as potential pathogens (5). Because avian erythrocytes are nucleated, we reasoned that this activity would be detectable in the steady state by TUNEL assay. Figure 2A shows that this is indeed the case. Most TIM4^++^ Kupffer cells contained multiple TUNEL-positive nuclei. To examine the phagocytic activity directly, *CSF1R*-mApple transgenic chickens were injected with 0.1 µm fluorescent latex beads. As shown in Figure 2B, after three hours these particles were co-located with TIM4. In this case, the *CSF1R*-mApple transgene was detected using anti-RFP antibody. The greater sensitivity of this detection revealed the presence of a population of stellate, *CSF1R*-mApple^+^, TIM4^−^ cells that were not actively phagocytic. The phagocytic activity of Kupffer cells was examined further following isolation *in vitro*. These isolated TIM4^+^ Kupffer cells exhibited strong phagocytic capability in uptake of fluorescence-labelled Zymosan particles (Figure 2C) apoptotic thymocytes (Figure 2D) and aged chicken red blood cells (Figure 2E).

**Figure 2.**
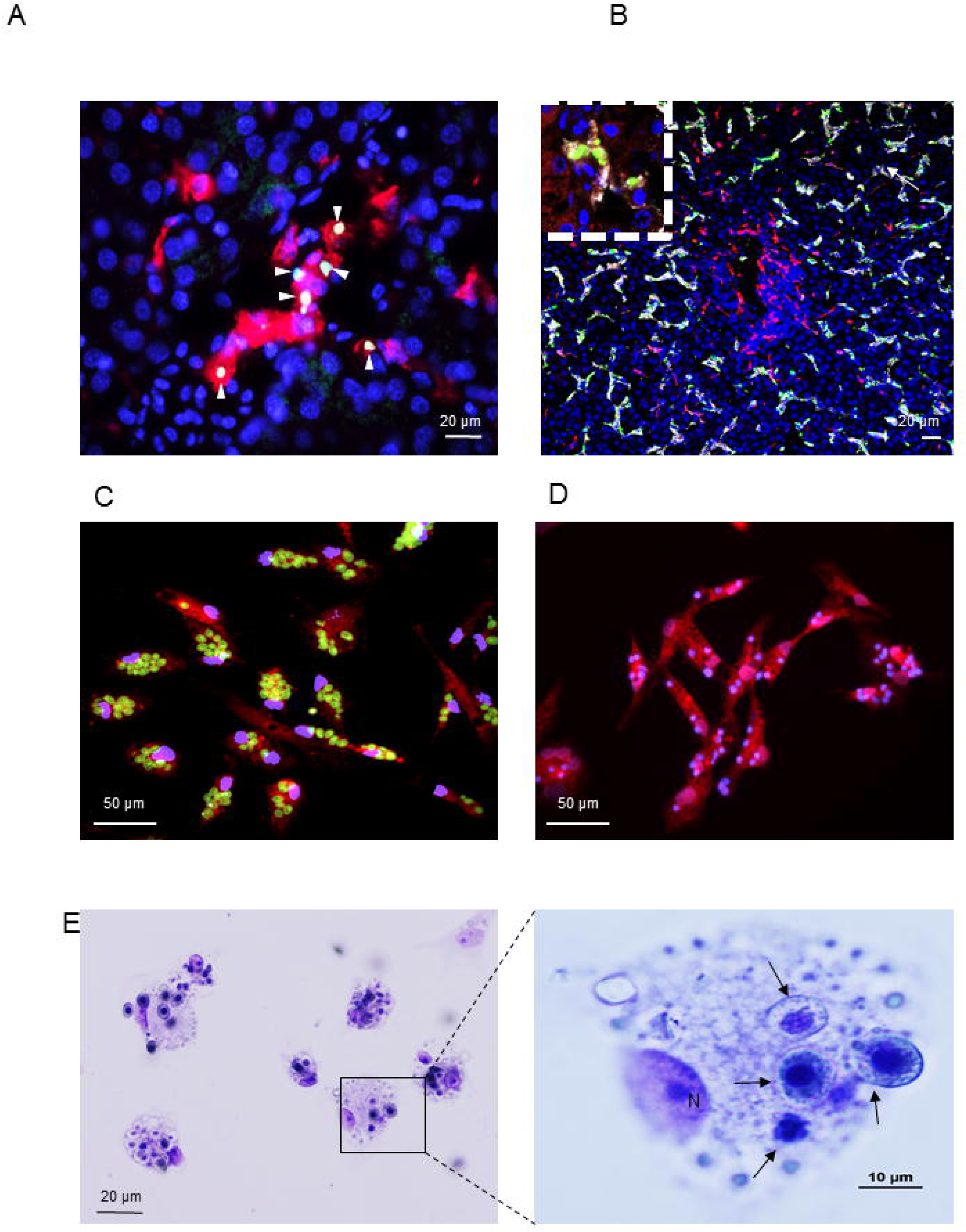
Phagocytic activity of TIM4 Kupffer cells. A). Clearance of apoptotic cells in liver. Cryostat sections from 4-week-old Novogen brown layers were analysed by TUNEL assay, the apoptotic cells in green and Kuffer cells identified by anti-TIM4 in red. B). *In vivo* study of Kupffer cell particle phagocytosis. Green fluorescent latex beads (0.1µ) were administrated to 4-week-old *CSF1R*-mApple transgenic birds via intravenous injection. Three hours later, the liver was harvested for cryosection. *CSF1R*-mApple cells were identified by localisation of mApple (red) and Kupffer cells with TIM4 (white). The nuclei were counter-stained with DAPI. Images produced by confocal microscopy using a Zeiss LSM 710. Inserted image shows the detail of TIM4^+^ Kupffer cell phagocytosis of green beads. C-E). *In vitro* analysis of Kupffer cell phagocytic activity. Kupffer cells were isolated from 4-week-old Novogen brown layers (n = 4) and cultured on a coverslip overnight. The adherent cells, mostly Kupffer cells, were mixed with zymosan fluorescent beads (C) or apoptotic thymocytes (D) at ratio 5 beads or apoptotic cells per cell. The cells were stained by anti-TIM4 antibody (red) and the nucleus counter-stained with DAPI (blue). Kupffer cell phagocytosis of chicken red blood cells was also analysed, as shown in light contrast image in (E).

In the bursa of Fabricius, a critical organ involved in avian B-cell development, only 1%-5% of B cells produced per day leave the organ, whereas many more cells undergo apoptosis *in situ* (29, 30). To examine the function of TIM4^+^ macrophages in clearance of apoptotic B cells, we double-stained bursal sections with anti-TIM4 and Bu-1. Apoptotic B cells (Bu-1^+^), with evidence of cell shrinkage and condensed chromatin, were apparently phagocytised by TIM4^+^ macrophages in the medulla and around the corticomedullary junction (Figure 3A). The phagocytosis of apoptotic B cells also occurred in embryonic bursa (Figure 3B). TUNEL assays confirmed the association of apoptotic B cells with TIM4^+^ macrophages in the bursa in both post-hatch birds (Figure 3C) and the embryo (Figure 3D).

**Figure 3.**
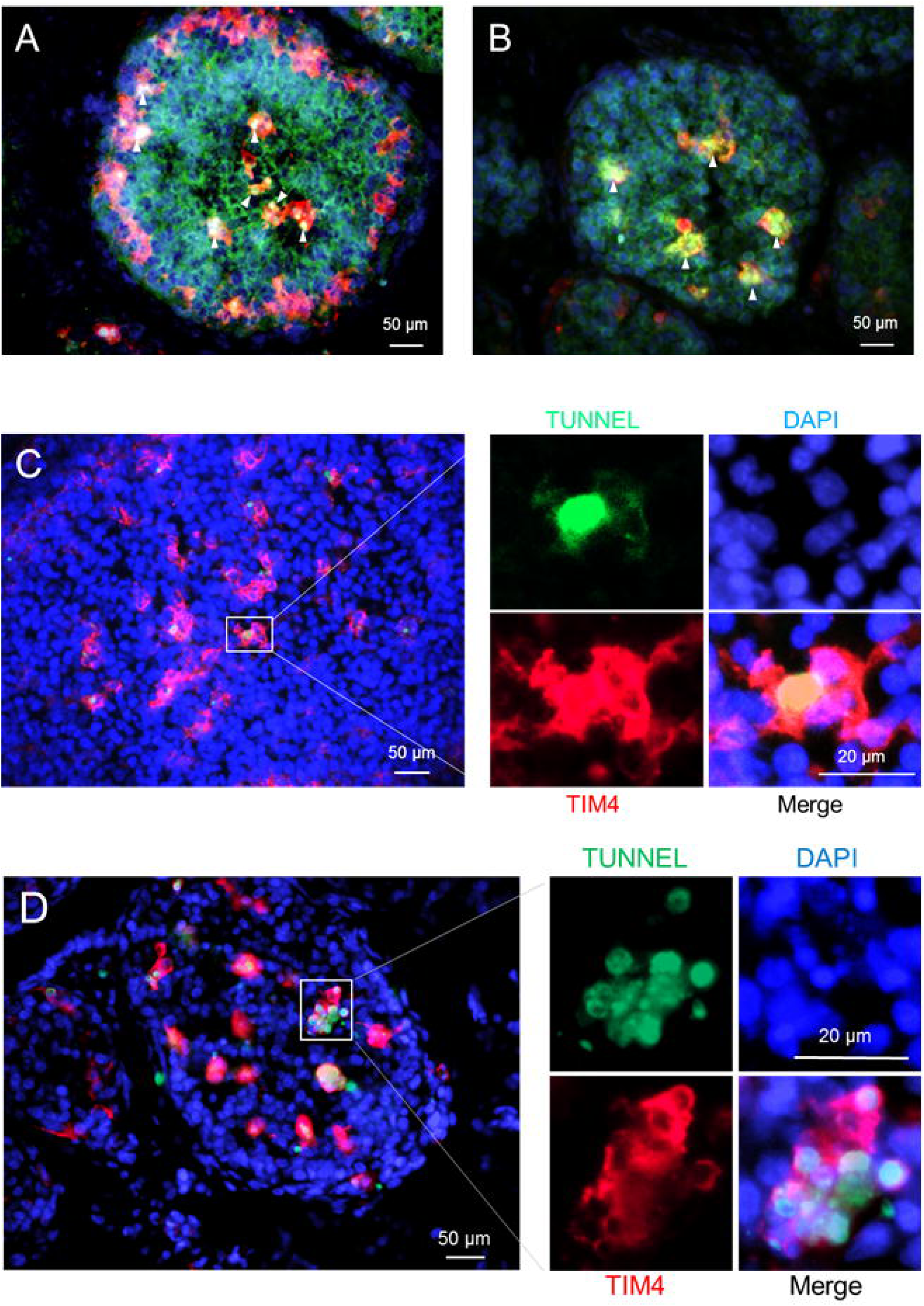
TIM4^+^ macrophage association with apoptotic B cells in bursa of Fabricius. Cryostat bursal sections (10 µm) from 3 week old birds (A) or embryos at day 18 (B) were co-stained for Bu-1 (green) and TIM4 (red) mAbs. The apoptotic B cell (Bu-1^+^) engulfed by TIM4 macrophages are indicated by arrows. Bursal sections from three week old birds (c) or embryos at day 18 (d) were also analysed by TUNEL to identify apoptotic cells (green) associated with TIM4+ macrophages (red). The nuclei were counterstained with DAPI (blue). The results show representative staining of three serial sections from at least three birds or embryos.

### The origins of TIM4^+^ macrophages in embryonic development

Macrophages first appear in the yolk sac prior to colonization of the embryonic body (15, 16, 18). An earlier study comparing mouse and chick, examined *TIMD4* mRNA in the chick semi-quantitatively at embryonic days (ED) 4-7, and inferred that *TIMD4* mRNA was likely expressed by yolk sac-derived macrophages. As noted in the introduction, this conclusion was supported by expression profiling of the chick embryo (19, 20). To confirm the location of TIM4 protein, we first examined yolk sac and embryos from embryonic day 3 (HH19) by whole mount immunohistochemistry (IHC). By ED3, blood island aggregates already contained numerous TIM4^+^ cells, and these were also clearly visible in vitelline blood vessels but had not yet appeared in the embryo (Figure 4A). At ED5, embryo-committed progenitors start to give rise to erythroid and monocyte progenitors (13). By this stage, TIM4^+^ cells were clearly detected and distributed in limb buds, the aorta-gonad-mesonephric (AGM) region, liver and ventricle (Figure 4C). TIM4^+^ cells isolated from disaggregated ED5.5 embryos and analysed by flow cytometry (Figure 4D) were CD45^+^ (>97%) and the majority also labelled with KUL01 (>67%).

**Figure 4.**
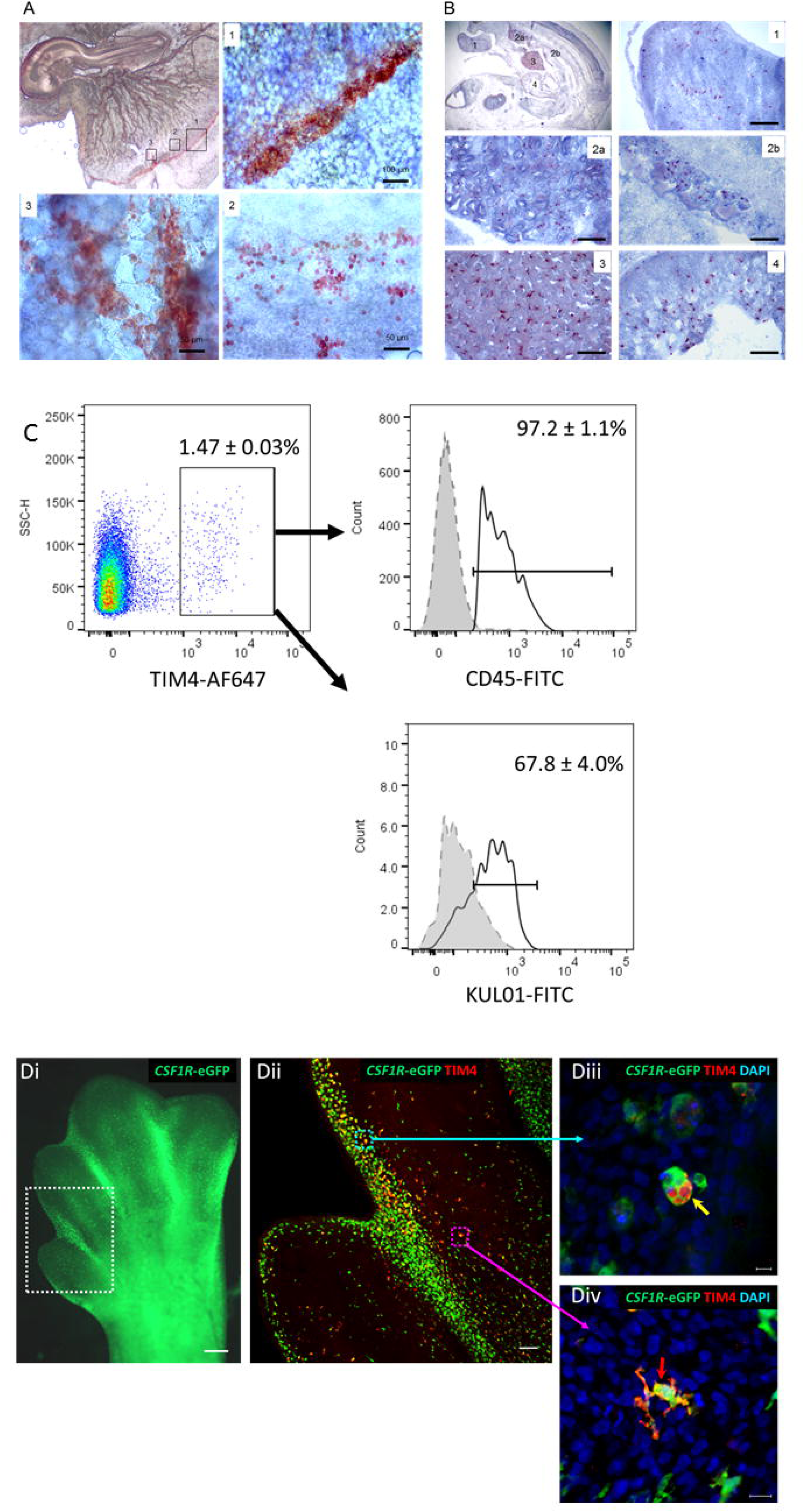
Localisation of the generation of TIM4+ macrophages in the yolk sac. A). Whole mount embryo staining. The whole ED 3 (HH 19) embryo was fixed and stained with anti-TIM4. Positive cells are stained with AEC (red) (10x). Higher magnification staining identifies positive cells in sinus terminals (1), 100 × magnification, blood island (2) and blood vessel (3) connected to sinus terminals, 200 × magnification. The results are representative of four embryos examined. B). Immunohistochemistry of midsagittal plane of ED5.5 embryo section. TIM4+ cells (AEC, red) and nucleus counter-stained by haematoxylin in blue. Whole embryo staining at top-left picture shows 10 × magnification. The detailed staining of limb bud (1), aorta-gonad-mesonephros region (2a and 2b), liver (3) and ventricle (4) is 200 × magnification with the scale bar 50 µm. Data represent one of three serial sections each from four embryos. C). Flow cytometric analysis of TIM4^+^ macrophages in ED5.5 embryos. Whole embryos (n=4) were dissociated by dispase/collagenase and resultant single cells double stained for TIM4 and either CD45 or KUL01 as indicated. Isotype control is indicated in grey-filled histograms. The numbers represent average percentages and error bars standard error of the mean. D). TIM4 staining on 7.5-8 day (HH33) limb bud from *CSF1R*-EGFP reporter chicks. (i) Distribution of EGFP+ macrophages in the chick limb bud. Scattered macrophages are found throughout the limb bud tissue and are especially concentrated in the interdigital region (boxed). Scale bar: 500 µm. (ii) 3D confocal reconstruction of a whole-mounted chick limb bud from the interdigital region shown in (i). This tile scan image represents a Z-stack containing 56 optical sections and shows EGFP+ macrophages and TIM4 staining. Scale bar = 50 µm. (iii) GFP+ macrophages in the interdigital region have a rounded morphology and contain TIM4+ phagolysosomes (yellow arrow), but no obvious surface staining. Scale bar = 10 µm. (iv) GFP+ macrophages outside the interdigital region have a ramified morphology, no obvious phagolysosomes and TIM4 staining is largely confined to the cell surface membrane Scale bar = 10 µm.

The same *CSF1R* regulatory elements used in *CSF1R-*mApple birds have also been used to drive expression of EGFP (14). In the *CSF1R*-EGFP chick, macrophages were visible throughout the body and were concentrated in areas of programmed cell death such as the interdigit regions of stage ED8 embryo leg buds (Figure 4D). TIM4 staining co-localised with these *CSF1R*-EGFP^+^ macrophages (Figure 4Dii). Interestingly, in macrophages associated with apoptotic cell remains in the interdigit region TIM4 staining appeared restricted to intracellular phagolysosomes, whereas in highly ramified macrophages outside this region TIM4 staining clearly delineated the plasma membrane (Figure 4Diii-iv). This pattern would be consistent with a role for TIM4 in apoptotic cell internalisation.

### Gene expression profiles of cells expressing TIM4

In mice, many tissue macrophage populations are maintained without major input from the blood monocyte pool (31). There is as yet no direct evidence that this is the case in birds, and our earlier data suggested that transplanted bone marrow progenitors can give rise to macrophages that populate most chick organs (17). Even in the mouse, organs such as skin and intestine are constantly replenished with macrophages derived from monocytes (15, 16). We therefore examined whether the extensive TIM4^+^ population in tissues might have precursors in blood in the *CSF1R*-mApple transgenic chicken by flow cytometric analysis (Figure 5A). As noted previously, the entire population of *CSF1R*-mApple^+^ blood monocytes expressed KUL01 and high levels of MHC II (14). Conversely, TIM4 staining clearly distinguished a subpopulation of positive cells, making up around 50% of the blood monocytes. These cells did not show any differential expression of class II MHC or KUL01 on flow cytometry. Combining the *CSF1R-*mApple transgene and TIM4 as markers, we separated the TIM4-positive and negative monocyte populations (Figure 5A) and assessed their gene expression by RNAseq. The primary data, comparing two pooled preparations of TIM4^+^ and TIM4^−^ monocytes separated as shown in Figure 5A, including relative expression ratios are provided in Table S1. *TIMD4* mRNA was enriched around 5-fold in the sorted TIM4^+^ population relative to TIM4^−^ moncytes, whereas *MRC1LB* (ENSGALT000000430910), which encodes KUL01 (26), was not different between the populations. The genes enriched within the TIM4^+^ cells include several globin subunits, and spectrin, suggesting that as in the liver, TIM4^+^ monocytes may be involved in uptake and destruction of senescent red blood cells. Other differences are discussed below.

**Figure 5.**
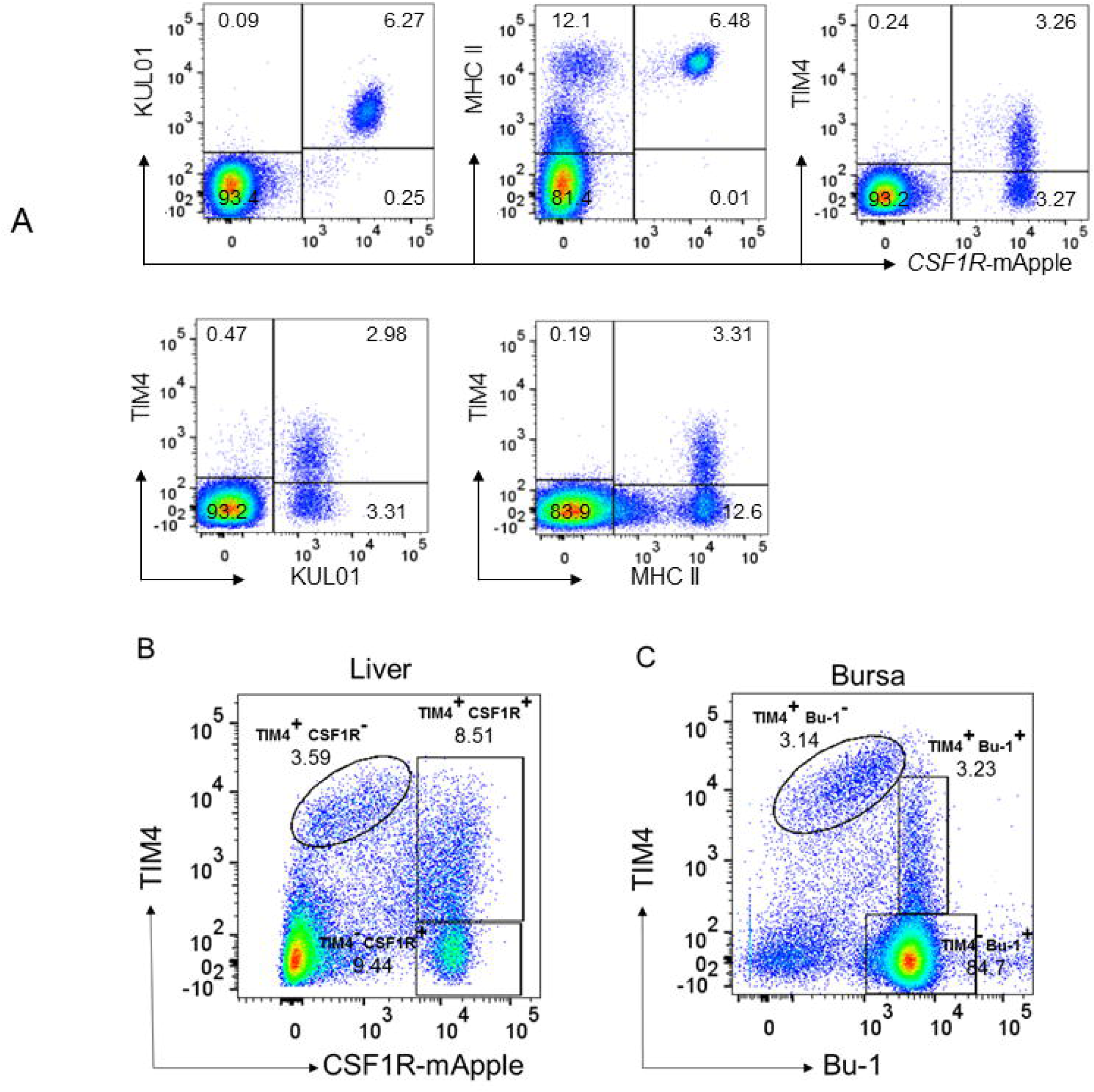
FACS separation of *CSF1R*-mApple+ and TIM4^+^ cells in blood, liver and bursa for RNAseq analysis A) Pooled blood leukocytes from *CSF1R-*mApple birds (n = 6) were labelled with anti-KUL01, MHCII or TIM4 as indicated. Note the uniform expression of KUL01 and MHCII on *CSF1R*-mApple+ cells, and heterogeneous expression of TIM4. B) Pooled non-parenchymal cells isolated from liver of *CSF1R*-mApple birds as described in Materials & Methods (n = 7). Note the separation of TIM4^+^*CSF1R*-mApple^−^ cells (Kupffer cells) from two populations *CSF1R*-mApple^+^ cells differing in expression of TIM4. C) Pooled isolated cell populations from bursa of Fabricius (n = 7) were double stained for TIM4 and Bu-1 as described in Materials and Methods. Note the presence of a population of TIM4^+^, Bu-1^−^ cells (macrophages) and two populations of Bu-1^+^ cells distinguished by TIM4 expression.

In single cell suspensions isolated from the liver, we identified three populations of cells separated by expression of the *CSF1R-*mApple and the level of expression of TIM4 (Figure 5B). Each population (TIM4^+^, *CSF1R-*mApple^−^; TIM4^+^, *CSF1R*-mApple^+^; and TIM4^−^, *CSF1R*-mApple^+^) was separated by FACS as shown in Figure 5B and mRNA expression profiles were assessed by RNAseq. Pairwise comparisons between the three populations are provided in Table S1. Surprisingly, the *CSF1R*-mApple^−^ populations expressed *CSF1R* mRNA at similar levels to the *CSF1R*-mApple^+^ cells, indicating that this transgene does not accurately report *CSF1R* transcription in the liver. *TIMD4* mRNA was 3-fold higher in the sorted TIM4^lo^, *CSF1R*-mApple^+^ cells relative to TIM4^−^, *CSF1R-*mApple^+^ cells, and a further 2.5-fold higher in the TIM4^hi^, *CSF1R*-mApple^−^ population.

Finally, in cells isolated from the bursa, we were not able to identify or recover sufficient cells for RNAseq profiling based upon the *CSF1R-*mApple transgene, despite the apparent prevalence of expressing cells in the tissue. Labelling with TIM4 and the B cell marker Bu-1 identified three populations which were separated by FACS using the gates shown in Figure 5C. mRNA expression profiles were again assessed by RNAseq. Pairwise comparisons of the three populations are shown in Table S1. Consistent with the isolation based upon TIM4, there was a hierarchy of *TIMD4* expression. *TIMD4* was just detected in the TIM4^−^ Bu-1+ population, was >10-fold higher in the TIM4^lo^, Bu-1^+^ population, and a further 10-fold higher in the TIM4^hi^ Bu-1^−^, population. As observed in the liver, despite the absence of detectable *CSF1R*-mApple, *CSF1R* mRNA was very highly-expressed in the TIM4^hi^ cells. The expression signatures of the liver and bursal TIM4^+^ populations are discussed further below.

### Transcriptional network analysis

To analyse the relationship of the separated cell populations to each other, and to identify transcripts that are strictly co-regulated with *TIMD4* and *CSF1R*, we utilised the network analysis tool *Graphia Pro* (www.kajeka.com), which was developed from BioLayout *Express*^3D^ (32). For this purpose, we included averaged expression from RNAseq datasets from hatchling spleen and bursa and from bone marrow-derived macrophages grown in CSF1 (12) as comparators. The spleen data set we analysed is derived from 18 male and female birds of diverse genetic background, in which *TIMD4* varied around 4-fold between individuals. For the current analysis, the male and female data were separately pooled and averaged. Figure 6 shows the sample to sample clustering. The analysis reveals the clear separation of blood monocytes, BMDM, bursa and spleen from all cells isolated from liver. The presumptive bursal B cells (Bu1^+^, TIM4^−^) most closely resemble the total spleen and bursa profiles and are clearly separated from macrophages. TIM4^+^ populations associated with their tissue/related cell type rather than with each other indicating that TIM4 expression is not, in itself, a differentiation marker. Gene-to-gene clustering identified sets of transcripts that were stringently co-expressed across the whole dataset. Annotated clusters with the average expression profile of each cluster are provided in Table S2. *TIMD4* was co-expressed with a small set of transcripts in Cluster 87, amongst which, only *CX3CR1* was expressed >50 TPM (*TIMD4* > 800 TPM). *CSF1R* was part of an even smaller cluster, Cluster 188, in which the only other robustly-expressed transcript is *CLCN5.* Clearly both *TIMD4* and *CSF1R* can be expressed by cells with very divergent cellular phenotypes and cannot be considered as markers of any particular cell type or lineage. Indeed, there were few clusters that were clearly associated with cell types or process. Not surprisingly, one of the largest clusters, Cluster 3, was clearly enriched for phagocyte-associated transcripts and components of the lysosome. A similar co-expression cluster was identified in mice (33); the highest average expression of amongst members of this cluster was in the chicken BMDM. Otherwise, the only other large expression clusters with evident cell type association were Cluster 7, which contains monocyte-enriched transcripts including *CD14, CCR2,, CSF3R, TLR2A* and transcription factors *CEBPB, NFE2L2, PRDM1* and *TFEC,* and Cluster 9, in which expression was highest in liver *CSF1R*-mApple^+^ cells. Consistent with the pairwise analysis above, this cluster contains Class II MHC and transcripts associated with dendritic cells (*BLB1, BLB2, CADM1, CIITA, CD74, CD86, FLT3, XCR1,* and transcription factors *IRF1, IRF5* and *IRF8*). The Kupffer cell marker, *MARCO*, was within a small cluster, Cluster 63, with GPR34 and *VSIG4*.

**Figure 6.**
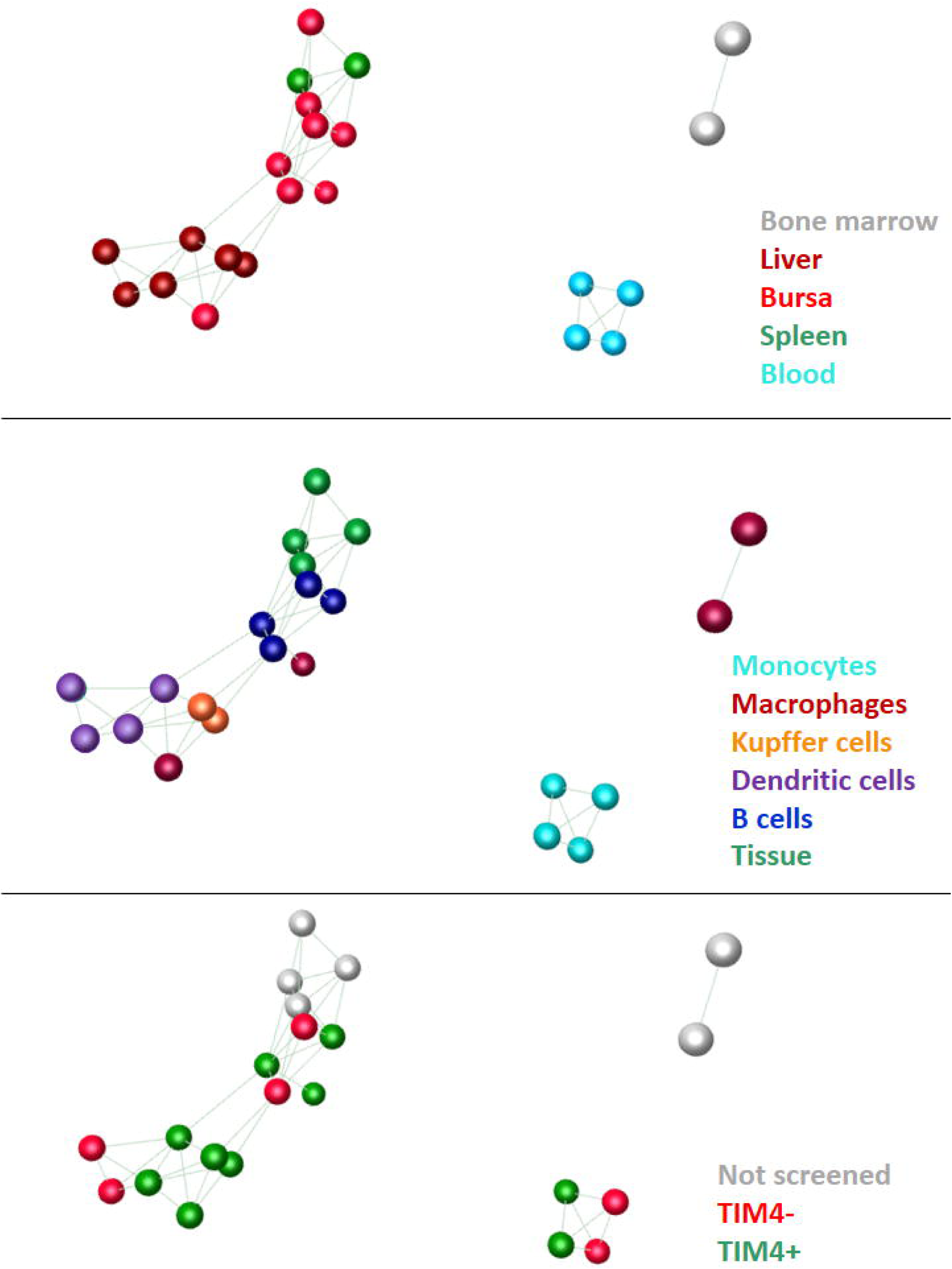
Sample to sample comparison of RNAseq profiles of isolated chicken cells and tissues RNAseq expression profiles for TIM4^+^ and TIM4^−^ blood monocytes (Figure 5A), liver populations separated based upon *CSF1R*-mApple and TIM4 (Figure 5B) and bursal populations separated based upon TIM4 and Bu-1 expression (Figure 5C) and were generated as described in Materials and Methods. For comparison, RNAseq expression data derived from female and male bone marrow derived macrophages (BMDM) and total spleen were pooled and averaged. Results from total bursa were derived from two male samples. All results were expressed as TPM. The dataset was filtered to remove transcripts where no sample reached a value of 1 TPM. Results were entered into Graphia and the data transposed to create a sample to sample matrix. Correlations with a Pearson correlation coefficient of ≥ 0.68 were entered into the analysis to allow all samples to be included. The three panels show the same data with node colours based upon tissue of origin (upper), cell type (middle) or TIM4 status (lower). In the middle panel, dendritic cells are the *CSF1R*-mApple^+^ cells from liver,

## Discussion

TIM4 in chickens, as in mammals, is a receptor expressed primarily by macrophages that binds to PS and most likely participates in the recognition and clearance of apoptotic cells. In this study, we used a novel anti-chicken TIM4 antibody to define the location of TIM4^+^ macrophages in adult chicken tissues and in developing chick embryos. In the embryo, macrophages are involved in extensive phagocytosis of dying cells, and the almost uniform co-expression of embryonic TIM4 with other macrophage markers (Figure 5) is consistent with that function. The data from post hatch birds (Figure 1 and 2) indicate that TIM4 is retained at the highest levels in a subset of tissue macrophages that appear to be associated specifically with the uptake of apoptotic cells and other particles.

We identified a subpopulation of blood monocytes in chickens that express detectable TIM4; around 50% of the total population (Figure 5A). *TIMD4* mRNA is not detected in monocytes in either mouse or human (34). Chicken blood monocytes uniformly expressed the *CSF1R*-mApple transgene, KUL01 (MRC1, CD206) and high levels of class II MHC. In mammals, two subpopulations of blood monocytes have been recognised and referred to as “classical” and “non-classical” (35). Aside from surface markers (Ly6C in mouse and CD16 in humans) the monocyte subpopulations in mammals differ especially in expression of the chemokine receptors, CCR2 and CX3CR1, which control their extravasation (36). The non-classical monocytes are derived from the classical monocytes, and their differentiation is controlled by CSF1 (31, 37). We used the combination of the *CSF1R-*mApple transgene and TIM4 to generate comparative gene expression profiles of chicken blood monocytes (Table S1). To our knowledge, this is the first such dataset generated. The cluster analysis in Table S2 identified a set of transcripts that was strongly enriched in chicken monocytes relative to other macrophage populations. In particular, *CSF3R* mRNA (encoding the G-CSF receptor) was very abundant. The ligand of this receptor in chickens was originally called myelomonocytic growth factor (MGF) (38). The level of expression of *CSF3R* suggests that it does indeed have a function in monocyte as well as granulocyte regulation in birds. The gene annotated as *CSF2RA* (GM-CSF receptor) was lowly-expressed, but a putative paralog (ENSGALT00000026942) was present at much higher levels in monocytes and on that basis, appears more likely to encode the CSF2 receptor. There are relatively few annotated monocyte surface markers in chick, but *MRC1LB* (KUL01), *CD14*, *MYD88*, *TLR4* and *TLR2A* were each highly-expressed and increased around 50% in TIM4^+^ cells. In common with mouse monocytes, the chicken monocytes expressed high levels of the chemokine receptors, *CCR2*, *CXCR4* and *CX3CR1.* The TIM4^+^ subpopulation had increased expression of *CX3CR1* in common with “non-classical” mouse monocytes. Other highly-expressed and monocyte-associated genes include those encoding a smorgasbord of transcription factors, notably *ATF4, CEBPA, CEBPB, CEBPD, EGR1, ELF1, ELF5, ETV6, FLI1, FOS, FOSB, FOSL2, HIC1, HIF1A, ILF3, IRF1, IRF2, IRF5, IRF8, JUN, JUND, KLF4, KLF6, MAF1, MAFB, MEF2D, MITF, NFE2L2, NFKB1, NFKB2, NR4A1, NR4A3, PPARD, RARA, REL, RUNX1, SPI1, STAT1, STAT3, STAT5B, STAT6,* and *TFEC*. Most of these factors have also been implicated in monocyte-macrophage differentiation in mice (39, 40). Most were marginally elevated in the TIM4^+^ chicken monocyte subset suggesting that these cells are part of a differentiation series rather than a distinct subset (Table S1). There are relatively few transcripts encoding specific functions that were both highly and selectively expressed in the TIM4^+^ monocytes. Most notable is around 2-fold increased expression of *ACVRL1, C1QA, C1QB, C1QC, CX3CR1, HAVCR1*, *ITGB5*, *LRP1*, *MARCO, MERTK, P2RY3, P2RY6, S1PR2, SMPD1* and *STAB1,* all of which have been implicated in apoptotic cell recognition and removal (41). The relative over-expression of the major alanine-rich C kinase substrates, *MARCKSL1* and *MARCKS* could also represent an adaptation for phagocytic activity (42). Accordingly, we suggest that chicken TIM4^+^ monocytes are adapted for apoptotic cell recognition. Given the extensive populations of TIM4^+^ macrophages in avian tissues, the circulating TIM4^+^ monocytes could provide progenitors to replace them, but equally, as in mammals, the tissue macrophages may self-renew and the TIM4^+^ monocytes may be the resident scavengers of blood.

The histological examination of the liver indicated that the TIM4^+^ positive phagocytes with the characteristic stellate morphology of Kupffer cells uniformly lacked expression of the *CSF1R-*mApple transgene (Figure 2) but they expressed *CSF1R* mRNA at the same level as the transgene-positive cells (Table S1). This finding appears at first glance to be distinct from mouse, in which a *CSF1R* reporter is expressed at high levels in all Kupffer cells (and indeed in all myeloid cells in the liver) (43). Kupffer cells in mice are rapidly depleted with anti-CSF1R antibody (37). They are the main site of clearance of CSF1 from the circulation (44) and respond to CSF1 with extensive cell proliferation (45, 46). So, mouse Kupffer cells clearly do express functional CSF1R. However, a modified transgene, which lacks a 150bp distal promoter element but contains the FIRE enhancer, like the chicken transgene, was active in monocytes and dendritic cells but undetectable in Kupffer cells or many other tissue macrophage populations (47, 48). There are regions of homology across avian species, aside from FIRE, that were not included in the avian *CSF1R* transgene (14). We infer that these are required for Kupffer cell *CSF1R* expression in chickens as in mice. Furthermore, most of the monocyte transcription factors noted above were down-regulated in Kupffer cells relative to the liver cells (Table S1) in which *CSF1R-*mApple was active; amongst these factors, FOS/JUN, IRF8, STAT1 and RUNX1 all bind to FIRE in mouse macrophages (40). Further consideration of *CSF1R* transcriptional regulation lies outside the focus of the current study. The key point is that the chicken *CSF1R* transgene we have produced provides a convenient marker that distinguishes monocytes and dendritic cells (discussed further below) from resident Kupffer cells.

The set of transcripts that was co-enriched with *TIMD4* in the TIM4^hi^, *CSF1R*-mApple^−^ Kupffer cell population (Table S1) includes transcripts expressed in hepatocytes (e.g. *ALB, TTR*) and endothelial cells (*CDH5, EDNRB, THBD*), which like the Kupffer cells, do not express the transgene. this probably reflect contamination, but might also involve apoptotic cell clearance. The Kupffer cell population may also be enriched for platelets. An unannotated gene (LOC101750889) of unknown function was highly-expressed and enriched in the Kupffer cells. The most-related protein in mouse and human genomes is the platelet membrane protein GP1BA, and *GP1BA* was itself highly-enriched in the TIM4^+^ population. Nevertheless, aside from *TIMD4* itself, relative to TIM4^lo^, *CSF1R-*mApple^+^ cells, the isolated TIM4^hi^ population selectively expressed numerous genes involved in the recognition and elimination of apoptotic cells, including *C1Q (A,B,C), CD36, CTSB, CTSD, CTSS, CX3CR1, DNASE2B, GPR34, LGALS1, LGALS3, MARCO, SCARB1, TGM2,* and *VSIG4* (highlighted in Table S1). Each of these genes is strongly Kupffer cell-enriched in mice (49, 50) and induced during embryonic liver differentiation (20). Notably, the scavenger receptor gene *MARCO* is almost exclusively expressed in the chick liver TIM4^+^ macrophage population (Table S1). One poorly-annotated liver-specific transcript that was even more highly-expressed than *MARCO, LOC101748207,* encodes a soluble scavenger receptor cysteine-rich domain-containing protein and may provide a novel Kupffer cell marker. The cluster analysis (Table S2) indicates that there are only few transcripts that share a transcription pattern with *MARCO,* of which only *VSIG4* and *GPR34* were highly-expressed. The underlying transcriptional regulation is also consistent with data from mice, in that genes encoding known regulators of apoptotic receptors in mice, *NR1H33* and *PPARG* (51) were also strongly-enriched in the chicken Kupffer cells. In mice, Kupffer cells are required for the elimination of senescent red blood cells and recycling of iron (52). Consistent with the conservation of this function and its regulation, the chick Kupffer cells were enriched for expression of the ferriportin gene, *SLC40A1*, the haeme transporter, *SLC48A1*, ferritin heavy chain, *FTH1* and haem oxygenase (*HMOX1*), and the transcription factor, *MAF,* which regulates expression of these genes (53) (Table S1). In mammals, CD163-mediated endocytosis of haemoglobin-haptoglobin complexes is a major pathway for iron uptake (54). However, *CD163*, and *CD163L1* were barely detected in any of the isolated chicken macrophage populations (Table S1).

A reciprocal set of genes was enriched in the TIM4^lo^, *CSF1R*-mApple^+^ population of liver macrophages relative to both Kupffer cells and the TIM4^−^ population (Table S1). The expression of the class II MHC genes *BLB1* and *BLB2*, and the class II invariant chain gene, *CD74*, was enriched more than 10-fold in this population relative to Kupffer cells (Table S1), along with the co-stimulators, *CD83* and *CD86* and monocyte/DC-associated transcription factors, *ATF3, BHLHE40, CIITA, FOS, IRF5, IRF8* and *NR4A3* and growth factor receptors, *CSF2RB* and *CSF3R.* Vu Manh *et al* (55) have previously isolated conventional dendritic cells (cDC) from chick spleen on the basis of high Class II MHC and the absence of KUL01. Like the chicken splenic cDC (55), the isolated TIM4^lo^ liver cells expressed low *MRC1LB* (encoding KUL01), and relative to Kupffer cells were also enriched for *BEND5, CADM1, CD40, CD86, CSF2RB, FLT3, LY75* (DEC205), *PLEKHA5, XCR1,* and *ZBTB46.* Table S1 identifies additional candidate markers for these cells: *LGALS2, LRRK1* and *P2RY6*. The data in Figure 2 and 3 demonstrate that these cells are not active phagocytes, but they are not deficient in lysosome-associated transcripts (e.g. *LAMP1, CTSB*) by contrast to isolated classical splenic DC in mice (33). Unlike the isolated chicken splenic DC, these cells also express *CSF1R* mRNA at the same level as Kupffer cells. DC-like cells have been identified in mouse liver (56), but they were a minor population, and largely concentrated under the capsule. In the chick, they are clearly considerably more numerous. Indeed, based upon the FACS profiles of isolated cells (Figure 5B), and localisation *in situ*, *CSF1R*-mApple^+^ liver DC are more abundant than Kupffer cells, and co-located in the sinusoids. Their gene expression profiles are related to blood monocytes, but monocytes lack *FLT3* mRNA. We speculate that in the absence of lymph nodes and limited lymphatics, the liver might be an important site of antigen recognition and presentation in birds.

The comparison of TIM4^+^ and TIM4^−^, *CSF1R*-mApple^+^ cells in the liver does not simply recapitulate the comparison in blood. The liver TIM4^+^ populations express *TIMD4* mRNA much more highly than TIM4^+^ monocytes. Again, the comparison is compromised somewhat by the apparent enrichment of hepatocyte-associated transcripts in the liver TIM4^+^ population. For that reason, the apparent expression of Kupffer cell-associated transcripts, such as *MARCO, VSIG4, C1QA, B, C* could represent a level of contamination with Kupffer cells. The more informative gene set, strongly-enriched in the TIM4^−^ population, includes monocyte-associated markers, *S100A8, CSF3R, TLR2A,* and *MRC1LB*, inflammatory cytokines (*IL1B, IL6*) and the stress-associated transcription factor, *NFE2L2.* A similar monocyte-like population has also been identified in mouse liver (56). However, the chick TIM4^−^ liver cells also express high levels of *FLT3, XCR1* and other DC markers noted above. It may be that they are a heterogeneous mix of cells, or an intermediate in differentiation from monocytes. This requires further investigation. Interestingly, the two *CSF1R*-mApple^+^ DC-like populations also express high levels of *SLC11A1*, also known as the natural resistance-associated macrophage protein 1 (*NRAMP1*), relative to much lower expression in TIM4^hi^ Kupffer cells and blood monocytes, BMDM and spleen (see Table S1 and Cluster 9 in Table S2). The chicken *SCL11A1* gene was previously shown to be expressed in liver, thymus and spleen. As in mice, *SCL11A1* polymorphism was associated with resistance to Salmonellosis (57). Consistent with that report, *SLC11A1* was undetectable in bursal cell populations (Table S1). The enrichment in the DC, which appear poorly endocytic, is somewhat paradoxical, since *Salmonella* is an intracellular pathogen, but suggests that the gene product may have a function in antigen presentation rather than control of intracellular pathogen replication. In overview, we have identified three populations of mononuclear phagocytes in the liver (Figure 5B), all of which express *CSF1R.* 20-30% of the isolated cells were TIM4^hi^ Kupffer cells. The gates in Figure 5B are somewhat arbitrary and isolation may not be quantitative. However, the relative abundance is consistent with the images in Figure 2. The remainder of liver mononuclear phagocytes express *CSF1R-*mApple and appear adapted for antigen presentation.

In the bursa, there were relatively few transcripts that were strongly-expressed and distinguish TIM4^+^ and TIM4^−^, Bu-1^+^ cells (Table S1). Those enriched in the TIM4^+^ population include *CSF1R,* transcription factors *SPIC*, *MAFA* and *MAFB*, and genes involved in apoptotic cell disposal including *CD274, CD244* and *VSIG4*. The levels of these transcripts were relatively low and we cannot eliminate a possible contribution from small numbers of contaminating macrophages or DC. Based upon high Bu-1 expression, and sample-to-sample clustering in Figure 6, both Bu-1^+^ populations appear to be mainly B cells, and indeed the TIM4^+^ and TIM4^−^ populations share similar high expression of the B cell receptor subunit, *CD79B*, the B cell kinase, *BTK,* class II MHC (*BLB1, BLB2*, *CD74*) and B cell transcription factors, *IKZF1*/*IKZF3*, *PAX5B*, *POU2F1*, and *TCF3.* Both populations are likely to actively proliferative, based upon their shared constitutive expression of cell cycle-associated genes, including regulators (*BUB1*, *CDC20, E2F*, *FOXM1*) which are almost absent in the TIM4^hi^, Bu-1^−^ cells. The level of TIM4, and of *TIMD4* mRNA, detected on these presumptive B cells was low, which may explain why they were not evident in sections of Bursa (Figures 1 and 2). The TIM4^hi^, Bu-1^−^ bursal cells, compared to TIM4^lo^, Bu-1^+^ cells, express high levels of *TIMD4* mRNA and share many enriched transcripts with Kupffer cells (e.g. *ACVRL1, C1QA, CIQB, C1QC, CX3CR1, FTH1, GPR34, HMOX1, LGALS3, LY86*, *MERTK, TGM2, SLC40A1, SPARC, STAB1*) consistent with adaptation for elimination of apoptotic cells and identity with the active phagocytes in Figure 3. However, they are distinct from Kupffer cells in expressing little *MARCO*, *VSIG4* and *CD36,* and in expressing very high levels of MHCII (*BLB1, BLB2, CD74*).

In conclusion, we have shown the specialised functional adaptation and transcriptional profile of liver TIM4^hi^ Kupffer cells in mammals is conserved in birds, and we have characterised a distinct population of phagocytes that express TIM4 in the bursa and which are also adapted to clear apoptotic cells. We have also identified functional diversity in chicken blood monocytes associated with expression of TIM4 and characterised a surprisingly abundant population of DC in the liver. Recent evidence in mice indicates that blood monocytes can, and do, differentiate into self-renewing Kupffer cells (58), even though this is not a major pathway in the steady state (31). Further studies will be required to determine whether *TIMD4*-expressing cells in the liver and bursa and elsewhere, and the distinct myeloid populations in liver, derive from monocytes or are entirely self-renewing. In this respect, our ability to generate cellular transplantation models, both *in ovo* and in hatchlings (17) and emerging capacity to generate knockouts (59) may make the chick a unique system for the study of macrophage and DC ontogeny.

## Acknowledgements

This paper is dedicated to the memory of our mentor, friend and colleague, Professor Pete Kaiser, who initiated this project. Pete passed away in July 2016. We would like to acknowledge Dr Taiana Pereira Da Costa at The Roslin Institute and R(D)SVS (current address: the Wildfowl & Wetlands Trust (WWT) in Slimbridge) for discussion on immunohistochemistry analysis and Dr Debiao Zhao for early guidance in embryo analysis.

## Footnote

This research was supported by the Biotechnology and Biological Sciences Research Council of the United Kingdom through Institute Strategic Programme Grants and project grant BB/M003094/1 to The Roslin Institute, and BB/H012559/1 to the former Institute for Animal Health (IAH). Support was also provided by the National Avian Research Facility funded by the Wellcome Trust with grant number 099164/Z/12/Z.

## References

1. Henson, P. M., and D. A. Hume. 2006. Apoptotic cell removal in development and tissue homeostasis. Trends Immunol 27: 244–250.

2. Poon, I. K., C. D. Lucas, A. G. Rossi, and K. S. Ravichandran. 2014. Apoptotic cell clearance: basic biology and therapeutic potential. Nat Rev Immunol 14: 166–180.

3. Bratton, D. L., V. A. Fadok, D. A. Richter, J. M. Kailey, L. A. Guthrie, and P. M. Henson. 1997. Appearance of phosphatidylserine on apoptotic cells requires calcium-mediated nonspecific flip-flop and is enhanced by loss of the aminophospholipid translocase. J Biol Chem 272: 26159–26165.

4. Miyanishi, M., K. Tada, M. Koike, Y. Uchiyama, T. Kitamura, and S. Nagata. 2007. Identification of Tim4 as a phosphatidylserine receptor. Nature 450: 435–439.

5. Theurl, I., I. Hilgendorf, M. Nairz, P. Tymoszuk, D. Haschka, M. Asshoff, S. He, L. M. Gerhardt, T. A. Holderried, M. Seifert, S. Sopper, A. M. Fenn, A. Anzai, S. Rattik, C. McAlpine, M. Theurl, P. Wieghofer, Y. Iwamoto, G. F. Weber, N. K. Harder, B. G. Chousterman, T. L. Arvedson, M. McKee, F. Wang, O. M. Lutz, E. Rezoagli, J. L. Babitt, L. Berra, M. Prinz, M. Nahrendorf, G. Weiss, R. Weissleder, H. Y. Lin, and F. K. Swirski. 2016. On-demand erythrocyte disposal and iron recycling requires transient macrophages in the liver. Nat Med 22: 945–951.

6. Wong, K., P. A. Valdez, C. Tan, S. Yeh, J. A. Hongo, and W. Ouyang. 2010. Phosphatidylserine receptor Tim-4 is essential for the maintenance of the homeostatic state of resident peritoneal macrophages. Proc Natl Acad Sci U S A 107: 8712–8717.

7. Albacker, L. A., P. Karisola, Y. J. Chang, S. E. Umetsu, M. Zhou, O. Akbari, N. Kobayashi, N. Baumgarth, G. J. Freeman, D. T. Umetsu, and R. H. DeKruyff. 2010. TIM-4, a receptor for phosphatidylserine, controls adaptive immunity by regulating the removal of antigen-specific T cells. J Immunol 185: 6839–6849.

8. Park, D., A. Hochreiter-Hufford, and K. S. Ravichandran. 2009. The phosphatidylserine receptor TIM-4 does not mediate direct signaling. Curr Biol 19: 346–351.

9. Amara, A., and J. Mercer. 2015. Viral apoptotic mimicry. Nat Rev Microbiol 13: 461–469.

10. Jemielity, S., J. J. Wang, Y. K. Chan, A. A. Ahmed, W. Li, S. Monahan, X. Bu, M. Farzan, G. J. Freeman, D. T. Umetsu, R. H. Dekruyff, and H. Choe. 2013. TIM-family proteins promote infection of multiple enveloped viruses through virion-associated phosphatidylserine. PLoS Pathog 9: e1003232.

11. Hu, T., Z. Wu, L. Vervelde, L. Rothwell, D. A. Hume, and P. Kaiser. 2016. Functional annotation of the T-cell immunoglobulin mucin family in birds. Immunology 148: 287–303.

12. Garceau, V., J. Smith, I. R. Paton, M. Davey, M. A. Fares, D. P. Sester, D. W. Burt, and D. Hume. 2010. Pivotal Advance: Avian colony-stimulating factor 1 (CSF-1), interleukin-34 (IL-34), and CSF-1 receptor genes and gene products. J Leukoc Biol 87: 753–764.

13. Garcia-Morales, C., L. Rothwell, L. Moffat, V. Garceau, A. Balic, H. M. Sang, P. Kaiser, and D. A. Hume. 2014. Production and characterisation of a monoclonal antibody that recognises the chicken CSF1 receptor and confirms that expression is restricted to macrophage-lineage cells. Dev Comp Immunol 42: 278–285.

14. Balic, A., C. Garcia-Morales, L. Vervelde, H. Gilhooley, A. Sherman, V. Garceau, M. W. Gutowska, D. W. Burt, P. Kaiser, D. A. Hume, and H. M. Sang. 2014. Visualisation of chicken macrophages using transgenic reporter genes: insights into the development of the avian macrophage lineage. Development 141: 3255–3265.

15. Epelman, S., K. J. Lavine, and G. J. Randolph. 2014. Origin and functions of tissue macrophages. Immunity 41: 21–35.

16. Ginhoux, F., and M. Guilliams. 2016. Tissue-Resident Macrophage Ontogeny and Homeostasis. Immunity 44: 439–449.

17. Garceau, V., A. Balic, C. Garcia-Morales, K. A. Sauter, M. J. McGrew, J. Smith, L. Vervelde, A. Sherman, T. E. Fuller, T. Oliphant, J. A. Shelley, R. Tiwari, T. L. Wilson, C. Chintoan-Uta, D. W. Burt, M. P. Stevens, H. M. Sang, and D. A. Hume. 2015. The development and maintenance of the mononuclear phagocyte system of the chick is controlled by signals from the macrophage colony-stimulating factor receptor. BMC Biol 13: 12.

18. Lichanska, A. M., and D. A. Hume. 2000. Origins and functions of phagocytes in the embryo. Exp Hematol 28: 601–611.

19. Lizio, M., R. Deviatiiarov, H. Nagai, L. Galan, E. Arner, M. Itoh, T. Lassmann, T. Kasukawa, A. Hasegawa, M. A. Ros, Y. Hayashizaki, P. Carninci, A. R. R. Forrest, H. Kawaji, O. Gusev, and G. Sheng. 2017. Systematic analysis of transcription start sites in avian development. PLoS Biol 15: e2002887.

20. Summers, K. M., and D. A. Hume. 2017. Identification of the macrophage-specific promoter signature in FANTOM5 mouse embryo developmental time course data. J Leukoc Biol.

21. Li, P. Z., J. Z. Li, M. Li, J. P. Gong, and K. He. 2014. An efficient method to isolate and culture mouse Kupffer cells. Immunol Lett 158: 52–56.

22. Wu, Z., L. Rothwell, T. Hu, and P. Kaiser. 2009. Chicken CD14, unlike mammalian CD14, is trans-membrane rather than GPI-anchored. Dev Comp Immunol 33: 97–104.

23. Bray, N. L., H. Pimentel, P. Melsted, and L. Pachter. 2016. Near-optimal probabilistic RNA-seq quantification. Nat Biotechnol 34: 525–527.

24. Soneson, C., M. I. Love, and M. D. Robinson. 2015. Differential analyses for RNA-seq: transcript-level estimates improve gene-level inferences. F1000Res 4: 1521.

25. Horn, F., A. M. Correa, N. L. Barbieri, S. Glodde, K. D. Weyrauch, B. Kaspers, D. Driemeier, C. Ewers, and L. H. Wieler. 2012. Infections with avian pathogenic and fecal Escherichia coli strains display similar lung histopathology and macrophage apoptosis. PLoS One 7: e41031.

26. Staines, K., L. G. Hunt, J. R. Young, and C. Butter. 2014. Evolution of an expanded mannose receptor gene family. PLoS One 9: e110330.

27. Gordon, S., and F. O. Martinez. 2010. Alternative activation of macrophages: mechanism and functions. Immunity 32: 593–604.

28. Houssaint, E., O. Lassila, and O. Vainio. 1989. Bu-1 antigen expression as a marker for B cell precursors in chicken embryos. Eur J Immunol 19: 239–243.

29. Lassila, O. 1989. Emigration of B cells from chicken bursa of Fabricius. Eur J Immunol 19: 955–958.

30. Motyka, B., and J. D. Reynolds. 1991. Apoptosis is associated with the extensive B cell death in the sheep ileal Peyer’s patch and the chicken bursa of Fabricius: a possible role in B cell selection. Eur J Immunol 21: 1951–1958.

31. Yona, S., K. W. Kim, Y. Wolf, A. Mildner, D. Varol, M. Breker, D. Strauss-Ayali, S. Viukov, M. Guilliams, A. Misharin, D. A. Hume, H. Perlman, B. Malissen, E. Zelzer, and S. Jung. 2013. Fate mapping reveals origins and dynamics of monocytes and tissue macrophages under homeostasis. Immunity 38: 79–91.

32. Theocharidis, A., S. van Dongen, A. J. Enright, and T. C. Freeman. 2009. Network visualization and analysis of gene expression data using BioLayout Express(3D). Nat Protoc 4: 1535–1550.

33. Hume, D. A., K. M. Summers, S. Raza, J. K. Baillie, and T. C. Freeman. 2010. Functional clustering and lineage markers: insights into cellular differentiation and gene function from large-scale microarray studies of purified primary cell populations. Genomics 95: 328–338.

34. Ingersoll, M. A., R. Spanbroek, C. Lottaz, E. L. Gautier, M. Frankenberger, R. Hoffmann, R. Lang, M. Haniffa, M. Collin, F. Tacke, A. J. Habenicht, L. Ziegler-Heitbrock, and G. J. Randolph. 2010. Comparison of gene expression profiles between human and mouse monocyte subsets. Blood 115: e10–19.

35. Ziegler-Heitbrock, L., P. Ancuta, S. Crowe, M. Dalod, V. Grau, D. N. Hart, P. J. Leenen, Y. J. Liu, G. MacPherson, G. J. Randolph, J. Scherberich, J. Schmitz, K. Shortman, S. Sozzani, H. Strobl, M. Zembala, J. M. Austyn, and M. B. Lutz. 2010. Nomenclature of monocytes and dendritic cells in blood. Blood 116: e74–80.

36. Auffray, C., M. H. Sieweke, and F. Geissmann. 2009. Blood monocytes: development, heterogeneity, and relationship with dendritic cells. Annu Rev Immunol 27: 669–692.

37. MacDonald, K. P., J. S. Palmer, S. Cronau, E. Seppanen, S. Olver, N. C. Raffelt, R. Kuns, R. Pettit, A. Clouston, B. Wainwright, D. Branstetter, J. Smith, R. J. Paxton, D. P. Cerretti, L. Bonham, G. R. Hill, and D. A. Hume. 2010. An antibody against the colony-stimulating factor 1 receptor depletes the resident subset of monocytes and tissue- and tumor-associated macrophages but does not inhibit inflammation. Blood 116: 3955–3963.

38. Leutz, A., K. Damm, E. Sterneck, E. Kowenz, S. Ness, R. Frank, H. Gausepohl, Y. C. Pan, J. Smart, M. Hayman, and et al. 1989. Molecular cloning of the chicken myelomonocytic growth factor (cMGF) reveals relationship to interleukin 6 and granulocyte colony stimulating factor. EMBO J 8: 175–181.

39. Monticelli, S., and G. Natoli. 2017. Transcriptional determination and functional specificity of myeloid cells: making sense of diversity. Nat Rev Immunol 17: 595–607.

40. Rojo, R., C. Pridans, D. Langlais, and D. A. Hume. 2017. Transcriptional mechanisms that control expression of the macrophage colony-stimulating factor receptor locus. Clin Sci (Lond) 131: 2161–2182.

41. Penberthy, K. K., and K. S. Ravichandran. 2016. Apoptotic cell recognition receptors and scavenger receptors. Immunol Rev 269: 44–59.

42. Sundaram, M., H. W. Cook, and D. M. Byers. 2004. The MARCKS family of phospholipid binding proteins: regulation of phospholipase D and other cellular components. Biochem Cell Biol 82: 191–200.

43. Sasmono, R. T., D. Oceandy, J. W. Pollard, W. Tong, P. Pavli, B. J. Wainwright, M. C. Ostrowski, S. R. Himes, and D. A. Hume. 2003. A macrophage colony-stimulating factor receptor-green fluorescent protein transgene is expressed throughout the mononuclear phagocyte system of the mouse. Blood 101: 1155–1163.

44. Bartocci, A., D. S. Mastrogiannis, G. Migliorati, R. J. Stockert, A. W. Wolkoff, and E. R. Stanley. 1987. Macrophages specifically regulate the concentration of their own growth factor in the circulation. Proc Natl Acad Sci U S A 84: 6179–6183.

45. Gow, D. J., K. A. Sauter, C. Pridans, L. Moffat, A. Sehgal, B. M. Stutchfield, S. Raza, P. M. Beard, Y. T. Tsai, G. Bainbridge, P. L. Boner, G. Fici, D. Garcia-Tapia, R. A. Martin, T. Oliphant, J. A. Shelly, R. Tiwari, T. L. Wilson, L. B. Smith, N. A. Mabbott, and D. A. Hume. 2014. Characterisation of a novel Fc conjugate of macrophage colony-stimulating factor. Mol Ther 22: 1580–1592.

46. Stutchfield, B. M., D. J. Antoine, A. C. Mackinnon, D. J. Gow, C. C. Bain, C. A. Hawley, M. J. Hughes, B. Francis, D. Wojtacha, T. Y. Man, J. W. Dear, L. R. Devey, A. M. Mowat, J. W. Pollard, B. K. Park, S. J. Jenkins, K. J. Simpson, D. A. Hume, S. J. Wigmore, and S. J. Forbes. 2015. CSF1 Restores Innate Immunity After Liver Injury in Mice and Serum Levels Indicate Outcomes of Patients With Acute Liver Failure. Gastroenterology 149: 1896–1909 e1814.

47. Ovchinnikov, D. A., C. E. DeBats, D. P. Sester, M. J. Sweet, and D. A. Hume. 2010. A conserved distal segment of the mouse CSF-1 receptor promoter is required for maximal expression of a reporter gene in macrophages and osteoclasts of transgenic mice. J Leukoc Biol 87: 815–822.

48. Sauter, K. A., C. Pridans, A. Sehgal, C. C. Bain, C. Scott, L. Moffat, R. Rojo, B. M. Stutchfield, C. L. Davies, D. S. Donaldson, K. Renault, B. W. McColl, A. M. Mowat, A. Serrels, M. C. Frame, N. A. Mabbott, and D. A. Hume. 2014. The MacBlue binary transgene (CSF1R-gal4VP16/UAS-ECFP) provides a novel marker for visualisation of subsets of monocytes, macrophages and dendritic cells and responsiveness to CSF1 administration. PLoS One 9: e105429.

49. Gautier, E. L., T. Shay, J. Miller, M. Greter, C. Jakubzick, S. Ivanov, J. Helft, A. Chow, K. G. Elpek, S. Gordonov, A. R. Mazloom, A. Ma’ayan, W. J. Chua, T. H. Hansen, S. J. Turley, M. Merad, G. J. Randolph, and C. Immunological Genome. 2012. Gene-expression profiles and transcriptional regulatory pathways that underlie the identity and diversity of mouse tissue macrophages. Nat Immunol 13: 1118–1128.

50. Mass, E., I. Ballesteros, M. Farlik, F. Halbritter, P. Gunther, L. Crozet, C. E. Jacome-Galarza, K. Handler, J. Klughammer, Y. Kobayashi, E. Gomez-Perdiguero, J. L. Schultze, M. Beyer, C. Bock, and F. Geissmann. 2016. Specification of tissue-resident macrophages during organogenesis. Science 353.

51. Roszer, T. 2017. Transcriptional control of apoptotic cell clearance by macrophage nuclear receptors. Apoptosis 22: 284–294.

52. Nairz, M., I. Theurl, F. K. Swirski, and G. Weiss. 2017. “Pumping iron”-how macrophages handle iron at the systemic, microenvironmental, and cellular levels. Pflugers Arch 469: 397–418.

53. Cairo, G., S. Recalcati, A. Mantovani, and M. Locati. 2011. Iron trafficking and metabolism in macrophages: contribution to the polarized phenotype. Trends Immunol 32: 241–247.

54. Graversen, J. H., M. Madsen, and S. K. Moestrup. 2002. CD163: a signal receptor scavenging haptoglobin-hemoglobin complexes from plasma. Int J Biochem Cell Biol 34: 309–314.

55. Vu Manh, T. P., H. Marty, P. Sibille, Y. Le Vern, B. Kaspers, M. Dalod, I. Schwartz-Cornil, and P. Quere. 2014. Existence of conventional dendritic cells in Gallus gallus revealed by comparative gene expression profiling. J Immunol 192: 4510–4517.

56. David, B. A., R. M. Rezende, M. M. Antunes, M. M. Santos, M. A. Freitas Lopes, A. B. Diniz, R. V. Sousa Pereira, S. C. Marchesi, D. M. Alvarenga, B. N. Nakagaki, A. M. Araujo, D. S. Dos Reis, R. M. Rocha, P. E. Marques, W. Y. Lee, J. Deniset, P. X. Liew, S. Rubino, L. Cox, V. Pinho, T. M. Cunha, G. R. Fernandes, A. G. Oliveira, M. M. Teixeira, P. Kubes, and G. B. Menezes. 2016. Combination of Mass Cytometry and Imaging Analysis Reveals Origin, Location, and Functional Repopulation of Liver Myeloid Cells in Mice. Gastroenterology 151: 1176–1191.

57. He, X.M., Fang, M.X., Zhang, Z.T., Hu, Y.S., Jia, X.Z., he, D.L., Liang, S.D., Nie, Q.H., and Zhang, X.Q. 2013. Characterization of chicken natural resistance-associated macrophage protein encoding genes (Nramp11 and Nramp2) and association with salmonellosis resistance. Genet. Mol. Res. 12: 618–630

58. Scott, C. L., F. Zheng, P. De Baetselier, L. Martens, Y. Saeys, S. De Prijck, S. Lippens, C. Abels, S. Schoonooghe, G. Raes, N. Devoogdt, B. N. Lambrecht, A. Beschin, and M. Guilliams. 2016. Bone marrow-derived monocytes give rise to self-renewing and fully differentiated Kupffer cells. Nat Commun 7: 10321.

59. Woodcock, M. E., A. Idoko-Akoh, and M. J. McGrew. 2017. Gene editing in birds takes flight. Mamm Genome 28: 315–323.

